# Chlorophyll fluorescence: How the quality of information about PAM instrument parameters may affect our research

**DOI:** 10.1101/2021.05.12.443801

**Authors:** Tim Nies, Yuxi Niu, Oliver Ebenhöh, Shizue Matsubara, Anna Matuszyńska

## Abstract

Chlorophyll *a* fluorescence is a powerful indicator of photosynthetic energy conversion in plants and photosynthetic microorganisms. One of the most widely used measurement techniques is Pulse Amplitude Modulation (PAM) fluorometry. Unfortunately, parameter settings of PAM instruments are often not completely described in scientific articles although their variations, however small these may seem, can influence measurements. We show the effects of parameter settings on PAM measurements. We first simulated fluorescence signals using a previously published computational model of photosynthesis. Then, we validated our findings experimentally. Our analysis demonstrates how the kinetics of non-photochemical quenching (NPQ) induction and relaxation are affected by different settings of PAM instrument parameters. Neglecting these parameters may mislead data interpretation and derived hypotheses, hamper independent validation of the results, and cause problems for mathematical formulation of underlying processes. Given the uncertainties inflicted by this neglect, we urge PAM users to provide detailed documentation of measurement protocols. Moreover, to ensure accessibility to the required information, we advocate minimum information standards that can serve both experimental and computational biologists in our efforts to advance system-wide understanding of biological processes. Such specification will enable launching a standardized database for plant and data science communities.

**Highlight:** PAM fluorometry measurement is sensitive to instrument settings and protocols. Yet, protocols are published incompletely. We urge to reach an agreement on minimal protocol information of PAM experiments to be shared publicly.

## 1 Introduction and the aim of this Communication

Oxygenic photosynthesis is one of the most essential processes on Earth. It drives the formation of oxygen and provides an energetic basis for carbon dioxide fixation. Due to its pivotal role in biomass production, the last decade has seen an increasing focus on engineering and manipulating photosynthesis in an attempt to improve plant productivity (Ort et al., 2015; Kromdijk et al., 2016; Kaiser et al., 2018; Flexas and Carriquí, 2020). Methods of quantifying the photosynthetic activity *in vivo*, in particular, the electron transport chain (Rochaix, 2011) allow inspection of dynamic changes in photosynthesis under variable environments. Foremost, measurement techniques based on chlorophyll *a* (Chl *a*) fluorescence have provided a broad range of information about reactions in photosystem II (PSII) and thylakoid membranes, leading to an upsurge in the understanding of photosynthesis (Kalaji et al., 2014, 2017).

A popular and important technique in photosynthesis research is Pulse Amplitude Modulation (PAM) Fluorometry (Schreiber et al., 1986). In combination with the saturation pulse method, it provides a minimally invasive system for the determination and quantification of the PSII activity (Schreiber, 2004). For detailed explanations of the method, its practical applications, and limitations, readers are directed to the excellent reviews published over the decades (Maxwell and Johnson, 2000; Baker, 2008; Murchie and Lawson, 2013). The basic principle of Chl *a* fluorescence and an example of induction measurement using PAM are shown in Fig. 1.

**Fig. 1:**
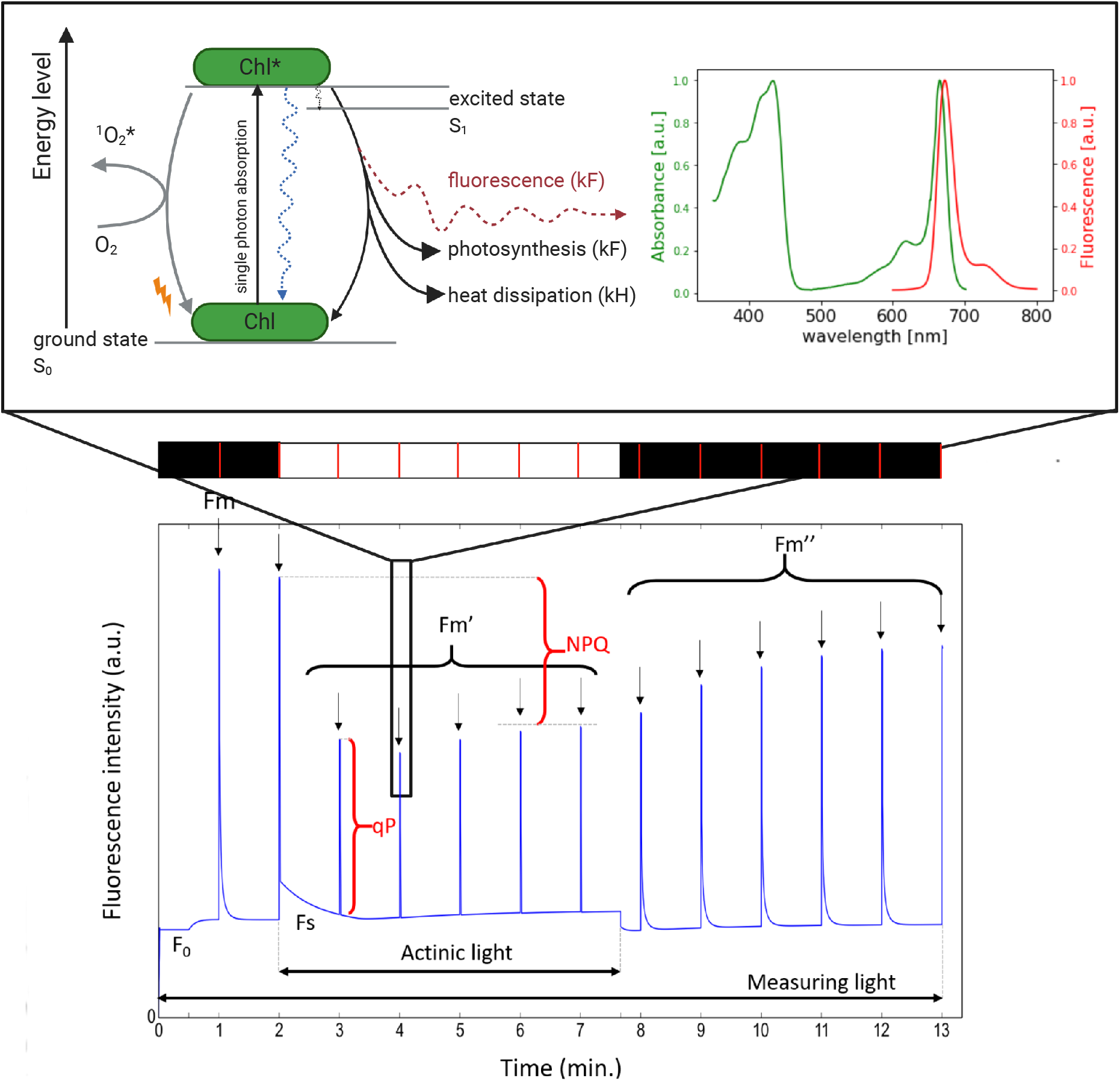
The basic principle of fluorescence and PAM induction measurement. In this protocol, a dark-adapted plant is exposed to actinic light (AL, white area shown in the light bar, L) followed by dark recovery (black area in L). Multiple saturating light flashes (saturation pulses, SP) are applied before, during and after the actinic illumination to measure the maximal fluorescence F_m_, F_m_’, F_m_”, respectively. Light energy absorption brings chlorophyll molecules to the first excited state (Si, see the simplified Jablonsky diagram in the upper panel on the left). For chlorophylls to return to the ground state (So), the absorbed energy can be used for charge separation and photosynthesis (photochemical quenching, qP), dissipated as heat (non-photochemical quenching, NPQ), or re-emitted as fluorescence. The fluorescence emission spectra are red-shifted from the absorption spectra (upper panel on the right) and can be detected using fluorometers with corresponding optical filters. The emitted fluorescence signal is recorded, as shown in the lower panel with key readouts. Based on Müller et al. (2001); Lichtman and Conchello (2006); Murchie and Lawson (2013). Created with BioRender.com

Non-photochemical quenching of Chl *a* fluorescence (NPQ) is one of the processes that can be quantified by analysing changes in fluorescence emission (Müller et al., 2001). The introduction of the PAM fluorometry opened up new opportunities for simple *in vivo* assessment of its dynamics. Under unfavourable conditions, NPQ serves as an important photoprotective mechanism, on one hand lowering the light use efficiency of photosynthesis, on the other protecting the photosynthetic apparatus from long term photo-damage (Ruban, 2016). The NPQ parameter is associated with the fraction of the light energy absorbed by PSII that is not used for photochemistry and is dissipated as heat.

Computational models serve as powerful tools for predicting systems’ responses to various changes and quantifying this effect. Their results help identify the reactions and mechanisms limiting photosynthetic productivity, to improve crop yield (Long et al., 2006; Zhao et al., 2020). Aspiring to make a similar impact through our research, our groups develop mathematical models of photosynthesis and actively search available fluorescence data to test computational models. Unfortunately, many articles presenting PAM fluorescence traces and data do not report detailed experimental protocols that were used to obtain these results. Such omission makes it challenging to reproduce fluorescence measurements by other groups but also *in silico*. This unintentional concealment of the experimental protocol is inevitably revealed once a computational approach is employed to replicate the experiment.

While simulating fluorescence traces using computational models we noted that our work required guessing some of the parameters used to conduct the experiment, as they were not explicitly stated in publications. This is not unexpected as incomplete reporting of experimental procedures has been identified as one of the factors responsible for the ‘‘reproducibility crisis” in science (Baker, 2016; National Academies of Sciences, Engineering, and Medicine, 2019; Jessop-Fabre and Sonnenschein, 2019). Most articles, in which PAM measurements were reported, included information about the type of fluorometer used (Sekulska-Nalewajko et al., 2019; Kalmatskaya et al., 2020) and the intensity of the saturation pulses (SP) (Vieira et al., 2013). Some, to our delight, attached the spectrum of actinic light used for fluorescence quenching analyses (Quero et al., 2020). But in many, values of the following four parameters were missing: i) the time interval (delay) between the determination of the maximum fluorescence (Fm) in darkness and switching on the actinic light (AL), ii) the intensity of the applied AL, iii) the time interval between the SPs, and iv) the duration of the SPs.

To systematically assess the effects of small variations in PAM parameters on the fluorescence traces, we simulated various PAM protocols using a mathematical model of NPQ published by our group (Matuszynska et al., 2016). We analysed the quantitative dependence of NPQ and PSII yield (Φ_PSII_) on the technical parameters that were mentioned above. Further, we validated the in *silico* findings by conducting two in vivo experiments in which the duration of the saturation pulse and the time point of switching on the actinic light were varied.

With this brief communication underlining the importance of full disclosure of PAM protocols in scientific publications, we hope to raise the awareness of authors, reviewers, and readers and thus to improve the knowledge transfer between the experimental and theoretical communities on plant physiology and photosynthesis. The findings presented here urges PAM users and the plant science community to consider facilitating broader exploitation of their data for modeling and meta-analysis studies while communicating their experimental procedures and results.

## 2 Materials and Methods

### 2.1 The model used for simulations

We use a mathematical model (Matuszyńska et al., 2016) to predict how changes in the values of four technical parameters of PAM measurement affect the fluorescence trace. For this, we simulate PAM experiments and systematically vary each of these parameters, quantifying the effect of each perturbation on NPQ and photosynthetic yield (Φ_PSII_). Table 1 contains descriptions of the standard variables derived from fluorescence signals, which are used to calculate NPQ and Φ_PSII_. The model comprises six ordinary differential equations (ODE) including a detailed description of NPQ. It has been parametrised for *Arabidopsis thaliana* and verified to accurately simulate fluorescence traces also for other higher plants. To reproduce the experimental results that are presented in this article we had to change three parameters. These are a coefficient of the light function transforming the photon flux density into the rate of excitation of PSII, the contribution of the protonated PsbS and zeaxanthin to the NPQ mechanism, and the proton leakage from the lumen in the stroma (a, *γ*_2_, k_leak_). All source code used to perform the presented analysis, together with the model implemented using the Python package `modelbase` developed by our group (van Aalst et al., 2021), can be downloaded from our git repository https://gitlab.com/qtb-hhu/fluopam.

**Table 1:** Parameters and descriptions of quantities derived from PAM fluorescence measurements in photosyhthetic organisms (Maxwell and Johnson, 2000; Murchie and Lawson, 2013).

| Parameter | Description |
| --- | --- |
| $F_m$ | maximal fluorescence in a dark-adapted state |
| $F_{m'}$ | maximal fluorescence in a light-exposed state |
| $F_o$ | minimal fluorescence in a dark-adapted state |
| $F_s$ | steady-state fluorescence level in a light-exposed state |
| NPQ | non-photochemical quenching $\frac{F_m - F_{m'}}{F_{m'}}$ |
| $\Phi_{\text{PSII}}$ | Quantum yield of photosystem II calculated as $\frac{F_{m'} - F_s}{F_{m'}}$ |

### 2.2 Standard PAM light induction protocol

We designed a PAM measurement that represents a generic experimental setup used with standard PAM fluorometers. Table 2 contains descriptions of the reference parameters. We have used 500 *μ*mol · s^-1^ m^-2^ as the default light intensity of AL. In the following analysis, we systematically varied the time point of switching on and off the AL, the intensity of the AL and SPs, and the duration and interval between SPs. We record the effect on the quenching capacity (NPQ) and photosynthetic yield (Φ_PSII_). Fig. 2 illustrates the basic idea behind our inquiries. We construct the light protocol (upper panel top) and solve the system of ODEs, from which we then calculate the fluorescence signal and plot it over time (upper panel bottom). From the fluorescence signal, we derive further the NPQ value (lower panel left) and quantum yield of photosynthesis (lower panel right).

**Table 2:**
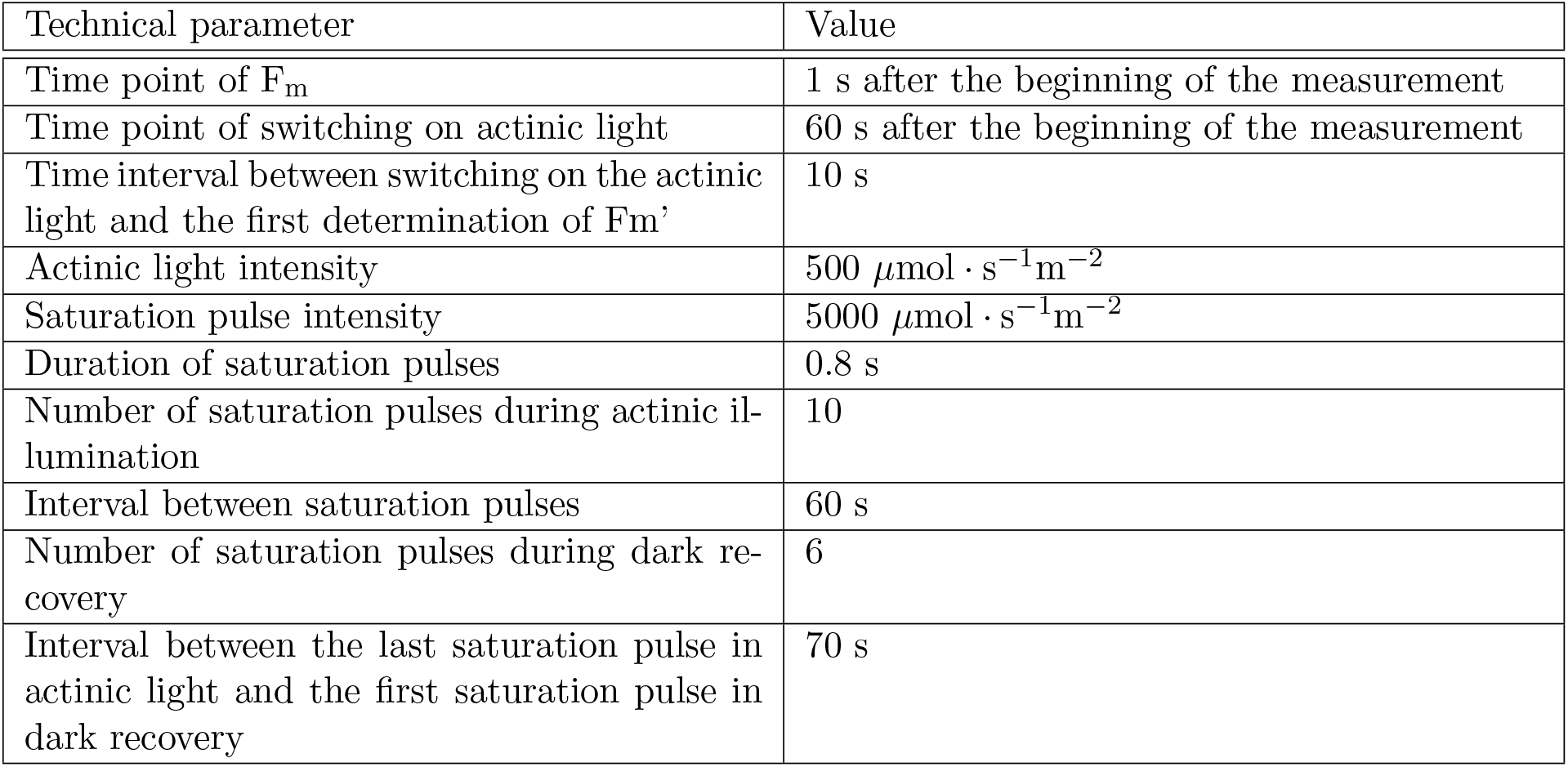
Parameters of the reference PAM protocol used for the simulations.

| Technical parameter | Value |
| --- | --- |
| Time point of $F_m$ | 1 s after the beginning of the measurement |
| Time point of switching on actinic light | 60 s after the beginning of the measurement |
| Time interval between switching on the actinic light and the first determination of $F_m'$ | 10 s |
| Actinic light intensity | $500 \mu\text{mol} \cdot \text{s}^{-1}\text{m}^{-2}$ |
| Saturation pulse intensity | $5000 \mu\text{mol} \cdot \text{s}^{-1}\text{m}^{-2}$ |
| Duration of saturation pulses | 0.8 s |
| Number of saturation pulses during actinic illumination | 10 |
| Interval between saturation pulses | 60 s |
| Number of saturation pulses during dark recovery | 6 |
| Interval between the last saturation pulse in actinic light and the first saturation pulse in dark recovery | 70 s |

**Fig. 2:**
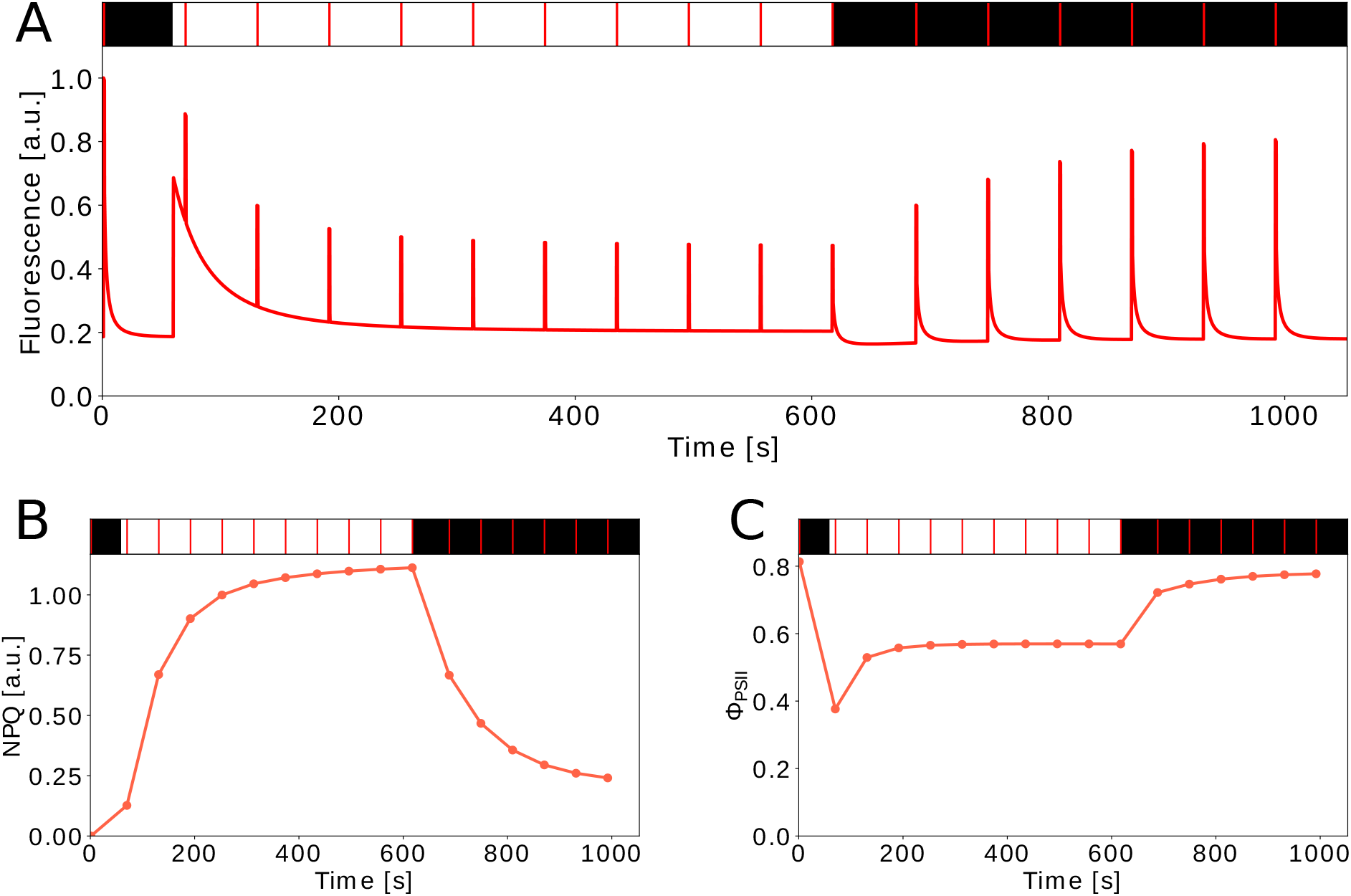
Example of the simulated PAM induction measurement with a saturation pulse protocol used in this work. Fig. 2A: the output of a typical simulated PAM fluorescence trace obtained for the reference parameters from Table 2. The dark/light/dark phases and times points of saturation pulses (in red) are indicated in the upper panel. Fig. 2B and 2C: the variables NPQ and Φ_PSII_, respectively, which are both derived from the simulated fluorescence.

### 2.3 Experimental methods

#### 2.3.1 Plant Material and Growth Conditions

Seeds of *Arabidopsis thaliana* (Columbia-0) were sown on moist commercial soil (Pikier, Balster Einheitserdewerk, Fröndenberg, Germany) and incubated at 4 °C in the dark. After three days they were transferred to a climate chamber with a 12 h/12 h light/dark photoperiod, 60% relative air humidity and 26 °C/20 °C day/night air temperature. The intensity of photosynthetically active radiation provided by fluorescent lamps (Fluora L58 W/77; Osram, Munich) was approx. 100 *μ*mol m^-2^s^-1^ at the plant height. Seedlings were transferred to pots (7 × 7 × 8 cm, one plant per pot) filled with soil (Lignostrat Dachgarten extensive, HAWITA, Vechta, Germany) on the 15^th^ day after sowing. Plants were watered from the bottom to keep soil moisture throughout the cultivation and during the experiments.

#### 2.3.2 PAM induction curve measurement

In the sixth week after sowing, Chl fluorescence measurements were performed in overnight dark-adapted plants using PAM-2500 (Walz, Effeltrich, Germany) equipped with leaf clip 2030-B. Before determination of the maximal PSII efficiency (F_v_/F_m_), 5 s of far-red light illumination (peak at 750 nm) was given to oxidize the electron transport chain and PSII. The intensity of red AL (peak at 630 nm) was set at approx. 457 *μ*mol m^-2^s^-1^. The default settings (10) were used for the intensity of both measuring light and SP. After 10 min of light induction, during which SP was applied every 60 s starting 1 s after the onset of AL illumination, AL was turned off and dark recovery was monitored for 13 min, during which seven SPs were applied with increasing time intervals as programmed in the protocol provided in the PamWin_3 software (Walz). In the experiment with varying time delay between the F_v_/F_m_ measurement and starting of the AL (10, 30, 40, 50, or 70 s), the width (duration) of SP was fixed at 800 ms (default). In the experiment with a varying width of SP (200, 400, 600, or 800 ms), the time delay was kept at 40 s (default). Four measurements were performed in four replicate plants (one measurement per plant) for each combination of the settings.

## 3 Results

### 3.1 Time Point of switching on and off the AL

In published protocols, we often encountered descriptions such as: “a dark-adapted plant has been exposed to a SP of light to measure the F_max_ and pulses were repeated every 30 seconds. After 12 minutes the light was switched off”. Such formulations, upon first reading, suggest that the plant has been exposed to AL immediately after the dark measurement of F_m_. However, by comparing our predicted fluorescence traces with the experimental data, we have noticed that in some of the experiments the initial “dark phase” must have continued after the first SP, sometimes even until the time point of the second SP. In Fig. 3A we illustrate the effect of the precise timing of switching on the AL on the predicted fluorescence. The time points of SPs are at seconds: 1 s, 70 s, 130.8 s, 191.6 s, 252.4 s, 313.2 s, 374 s, 434.8 s, 495.6 s, 556.4 s, 617.2 s, 688 s, 748.8 s, 809.6 s, 870.4 s, 931.2 s, 992 s. Naturally, the timing of switching on the AL has implications for the NPQ induction while the subsequent dark relaxation is not affected (Fig. 3B). The time course of F_s_ quenching under the AL is unaffected although the F_s_ level at a given time point changes (Fig. 3A). As expected, the longer the dark phase between the first two SPs, the lower the initial NPQ determined by the second SP applied. These simulations demonstrate that, without precise knowledge of this parameter, a rigorous model interpretation of the NPQ induction kinetics is difficult.

**Fig. 3:**
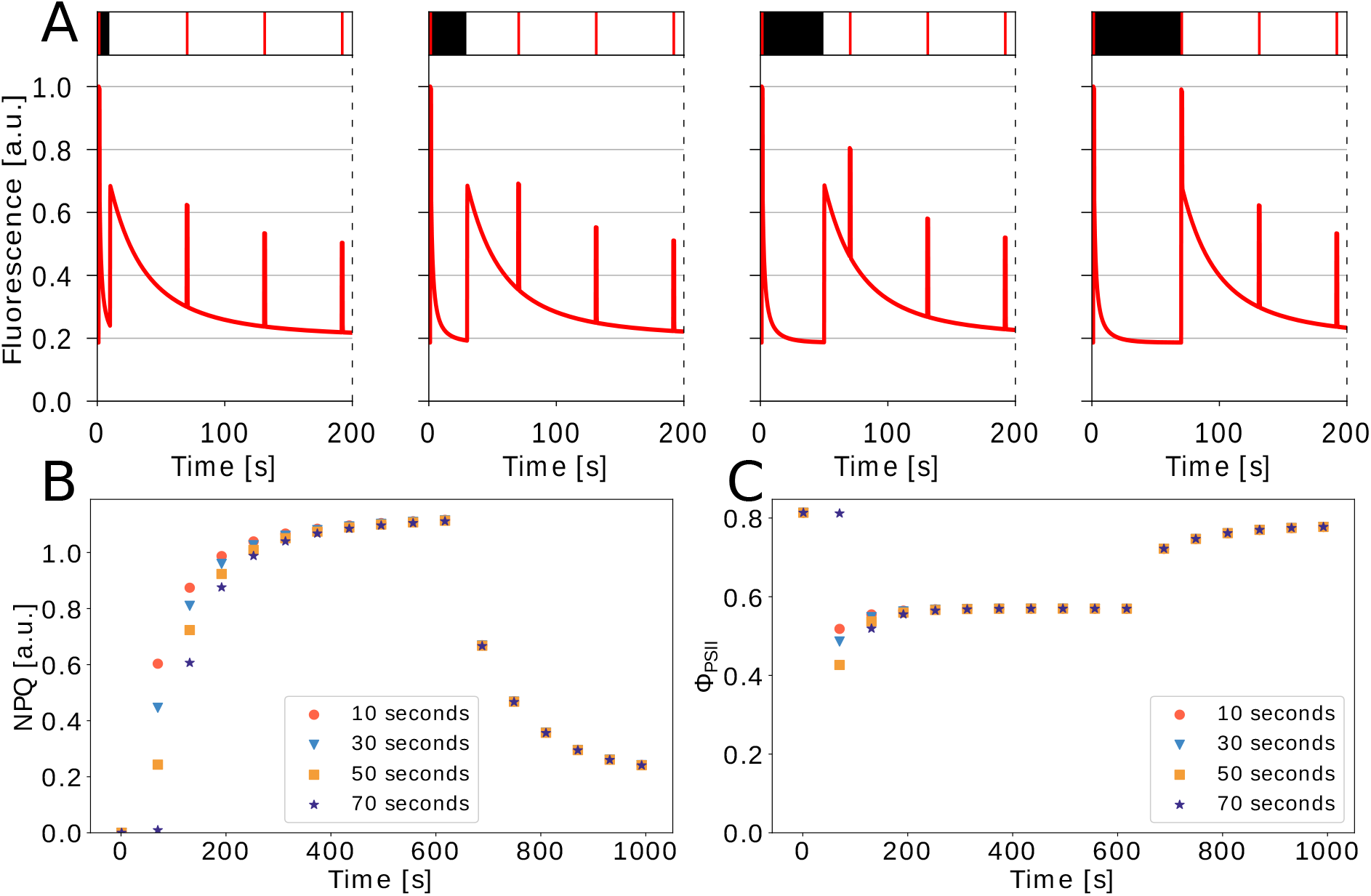
Variation of the time point of switching on the AL. Fig 3A: excerpts of full fluorescence traces. The time point of switching on the AL is varied while the time points of applying SP are kept unchanged. From left to right: 10 s, 30 s, 50 s, and 70 s after the first SP to determine F_m_. The vertical dashed lines indicate that the fluorescence traces continue. Fig. 3B: derived NPQ values. Fig. 3C: yield of photosystem II (Φ_PSII_).

Likewise, the exact time of switching off the AL is also not described in publications. In Fig. 4 we have additionally varied the time point of switching off the AL in a similar manner as done above for switching on the AL while maintaining the overall duration of the AL. In the upper panel the four light-to-dark transition phases are plotted and in the lower panels the derived NPQ and ΦPSII. Additionally to the apparently altered induction kinetics of NPQ observed before, now significant variations in the dark relaxation kinetics can be clearly seen in Fig. 4B.

**Fig. 4:**
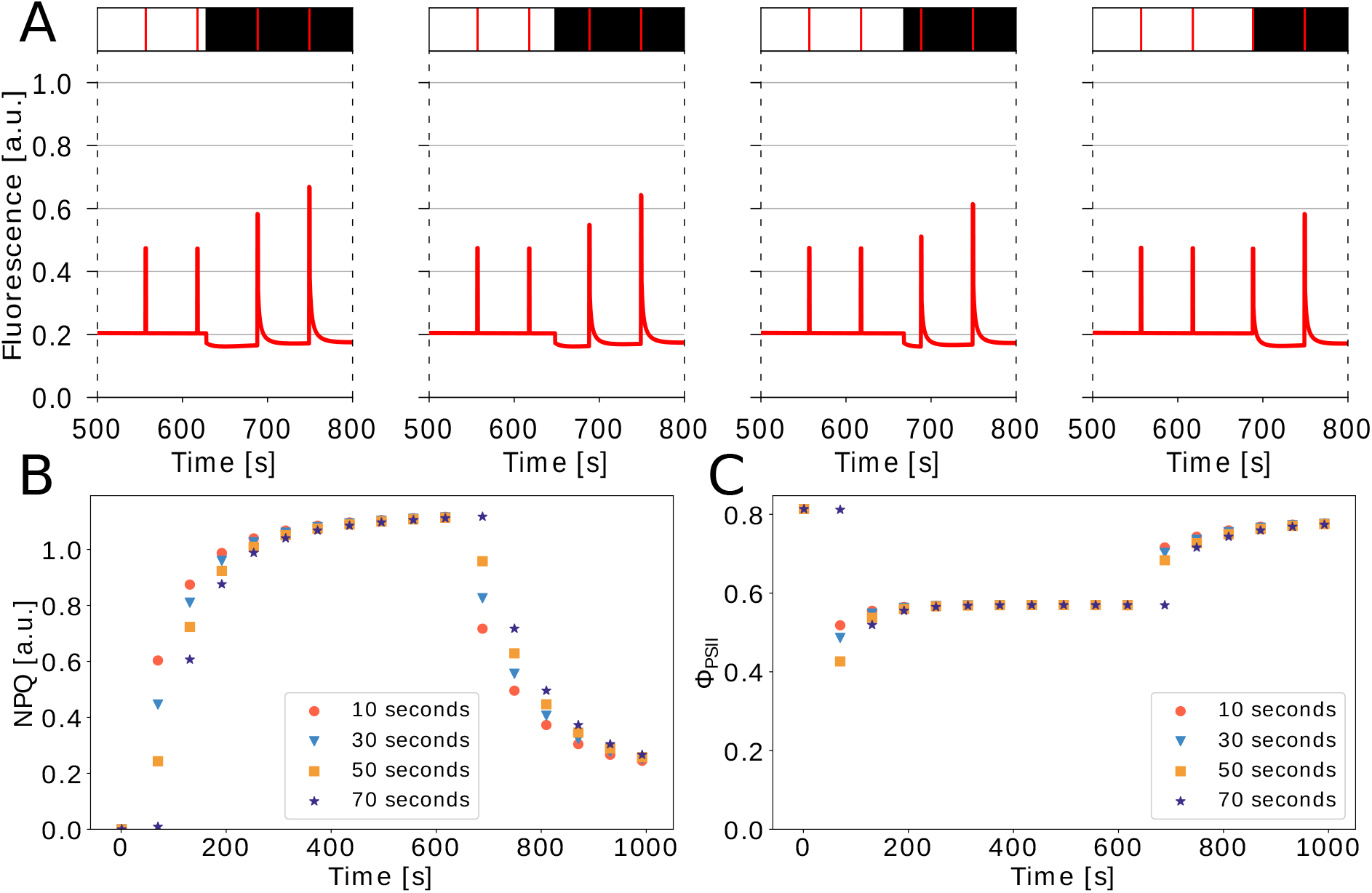
Variation of the time point of switching on and off the AL. Fig. 4A: excerpts of full fluorescence traces. The time point of switching on and off the AL is varied while the time points of applying SP are kept unchanged. From left to right 10 s, 30 s, 50 s, and 70 s after first SP for Fm and a corresponding shift of the last SP in the AL, respectively. Vertical dashed lines indicate that the fluorescence traces continue. Fig. 4B: derived NPQ values. Fig. 4C: yield of photosystem II (Φ_PSII_).

### 3.2 AL and SP intensity

The precise definition of the AL is essential for an accurate interpretation of raw fluorescence traces, as well as other dynamic variables derived from them. Commonly used expressions such as ‘moderate light’ or ‘low light intensity’ are not informative. Light activation depends on certain physical, biochemical, and structural parameters that vary between photosynthetic organisms depending on e.g. the chlorophyll content, the threedimensional structure of the chloroplast, and the composition of the thylakoid membrane. In our model, these differences are accounted for in the light activation function. In this analysis, we systematically increased the AL from intensity 100 to 1000 *μ*mol · s^-1^m^-2^ to examine the effect on the derived steady-state NPQ value. As shown in Fig. 5, the intensity of AL influences the steady-state NPQ level significantly, in this example up to 350 *μ*mol · s^-1^m^-2^ where it reaches saturation.

**Fig. 5:**
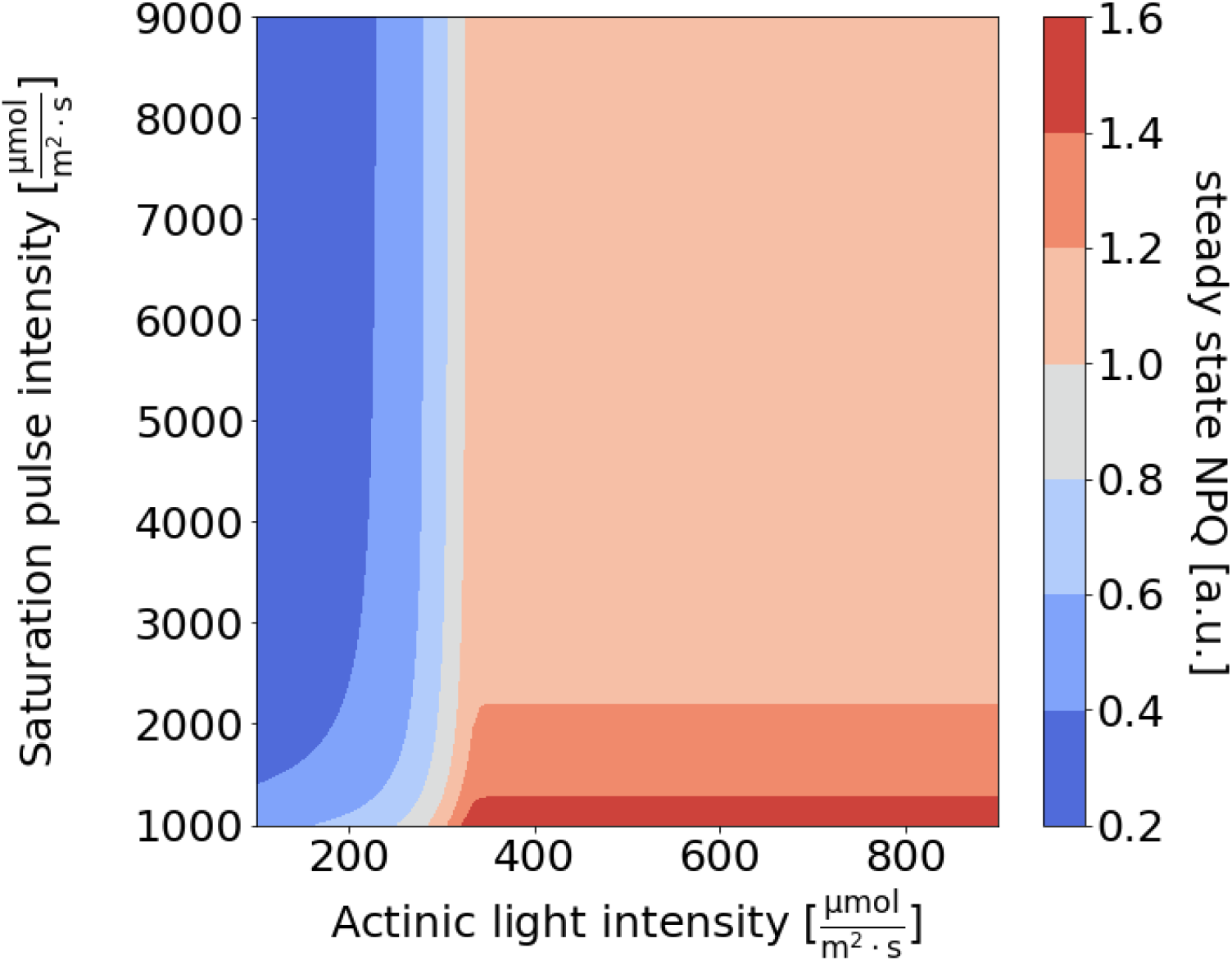
Steady-state NPQ values (derived at the last SP in the AL environment) simulated for various combinations of AL and SP intensities.

Besides the intensity of AL, the intensity of the SPs also plays a crucial role in the reproducibility of PAM experiments. To study possible consequences of different SP intensities, we altered the values between 1000 and 9000 *μ*mol · s^-1^ m^-2^. The calculated steady-state NPQ values are higher for SP intensities below 3000 *μ*mol · s^-1^m^-2^, suggesting that only for intensities over 4000 *μ*mol · s^-1^m^-2^ the SP is really saturating. This is an interesting finding of our analysis, where with the simulations we could identify the theoretical minimal intensity of the SP which, in this example, was about 10-fold higher than the lowest AL intensity to induce the maximal NPQ (Fig. 5).

### 3.3 Interval between SPs

Next, we investigated the effect of the time interval between the SPs. From our experience, the value of this parameter is explicitly mentioned in far more articles than for instance the time of switching on the AL, indicating the importance given to this parameter. Fig. 6 shows derived NPQ and Φ_PSII_ values from the PAM protocols with varying intervals between the SPs. Based on the results shown in Fig. 6 one could conclude that this technical parameter, if it is within the range typically used by many groups in laboratory experiments (30, 60, 120 s), may not alter the induction and relaxation kinetics or the steady-state level of NPQ. However, when we repeated the analysis for a lower AL intensity of 100 *μ*mol · s^-1^m^-2^, a tendency of increased NPQ by short-interval SPs could be observed under the AL as well as in the subsequent darkness (Fig. 6C).

**Fig. 6:**
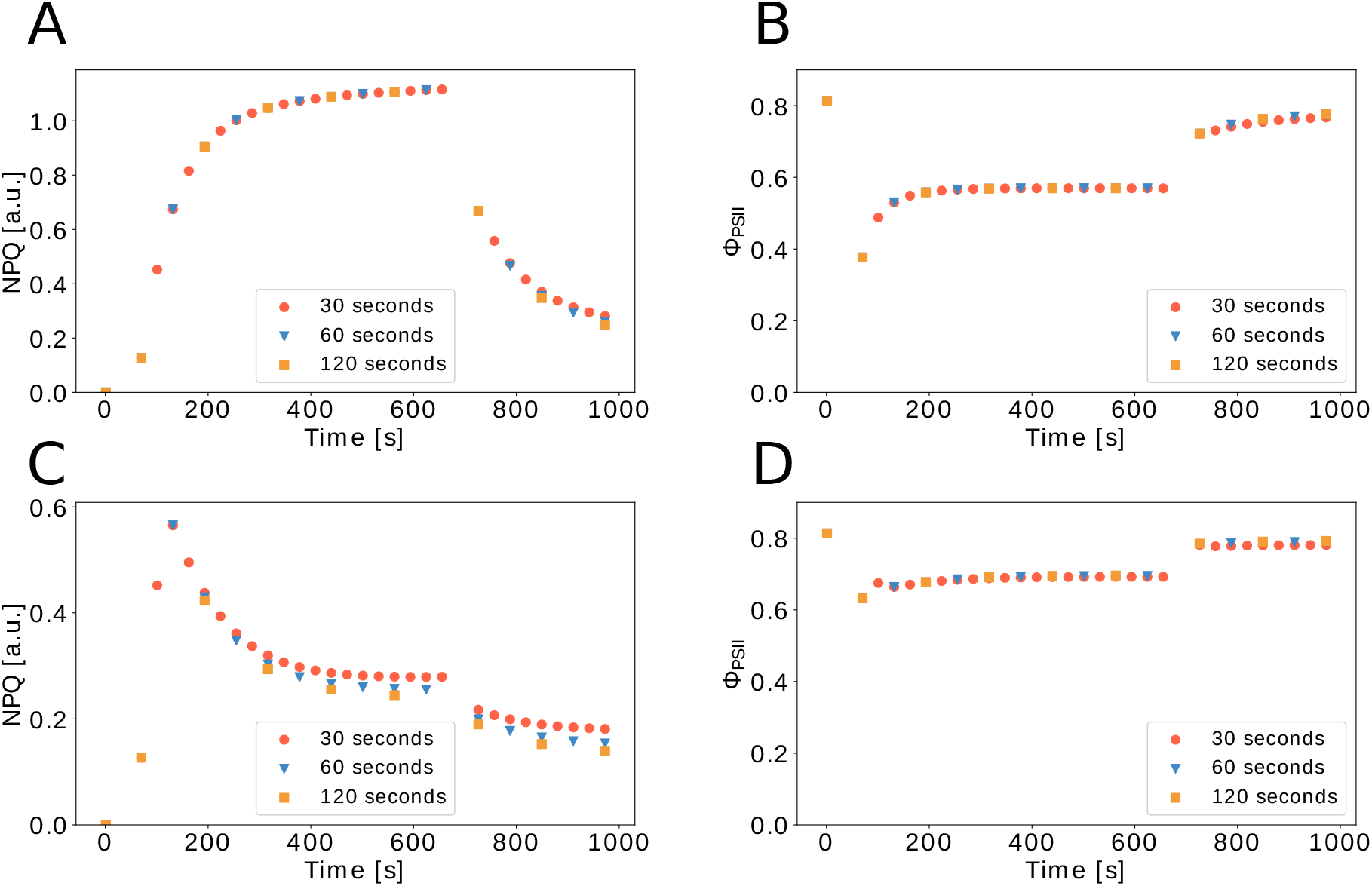
NPQ and Φ_PSII_ values simulated for the standard PAM induction protocol with varying time intervals between two consecutive SPs. 6A and 6A: with AL intensity of 500 *μ*mol · s^-1^m^-2^. 6C and 6D: with AL intensity of 100 *μ*mol · s^-1^m^-2^

### 3.4 Duration of SP

Lastly, we investigated the effect of the SP duration. It is often assumed that the SPs of light have no lasting effect on the photosynthetic system, as long as they are ‘short’ (Schreiber, 2004). To verify this claim, we examined the effect of the SP length from 0.2 s to 2.8 s (Fig. 7). Our simulation confirmed no effect of the duration of the SP at 5000 *μ*mol · s^-1^m^-2^ and all investigated intervals on NPQ under the used AL intensity. Interestingly, this technical parameter was among those that are most regularly and explicitly mentioned in the method sections. Despite the special attention and notes, however, there seems to be only a minor immediate effect of altering this parameter, as indicated by our analysis in Fig. 7.

**Fig. 7:**
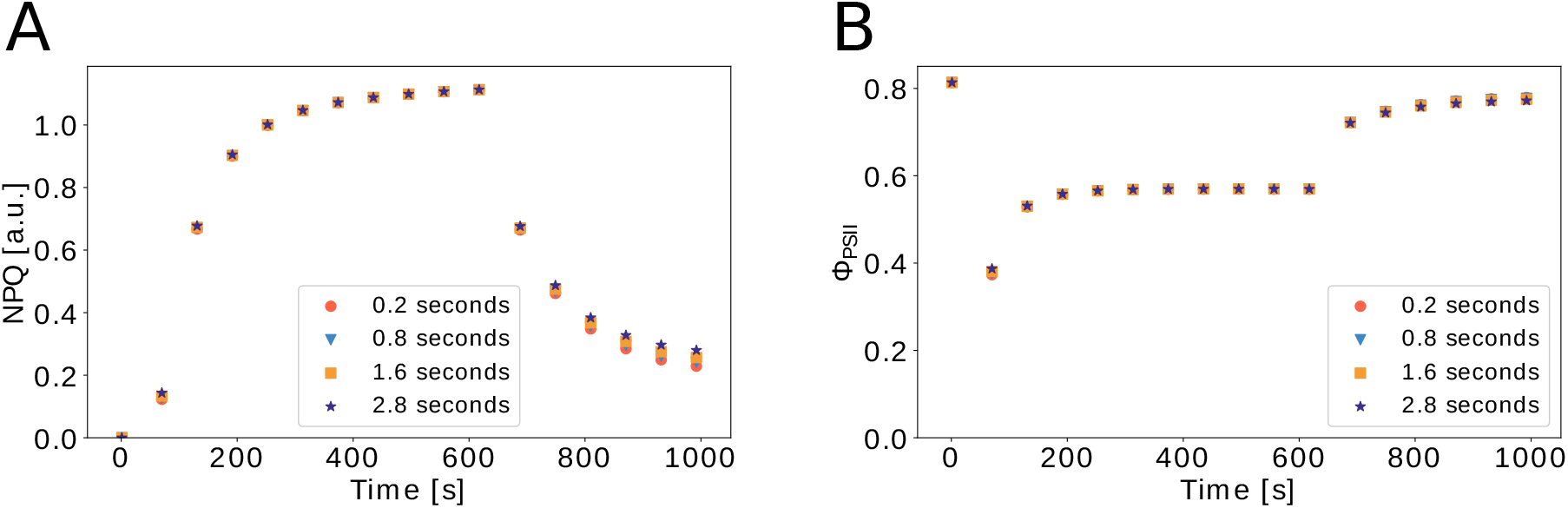
NPQ and Φ_PSII_ values simulated for the standard PAM induction protocol with varying length of the saturation light pulses.

### 3.5 Model validation

To validate and support our *in silico* analyses we conducted two experiments using plants grown as described in the Method section. The first experiment investigates the impact of the duration of the SP (referred to as SP experiment). The second one focuses on the time point of switching on the AL with the first SP in light-triggered after 1 s (referred to as Delay experiment). As Figs. 8 and 9 show, our simulations are qualitatively in good agreement with the experimental data of NPQ and Φ_PSII_ in both experiments.

**Fig. 8:**
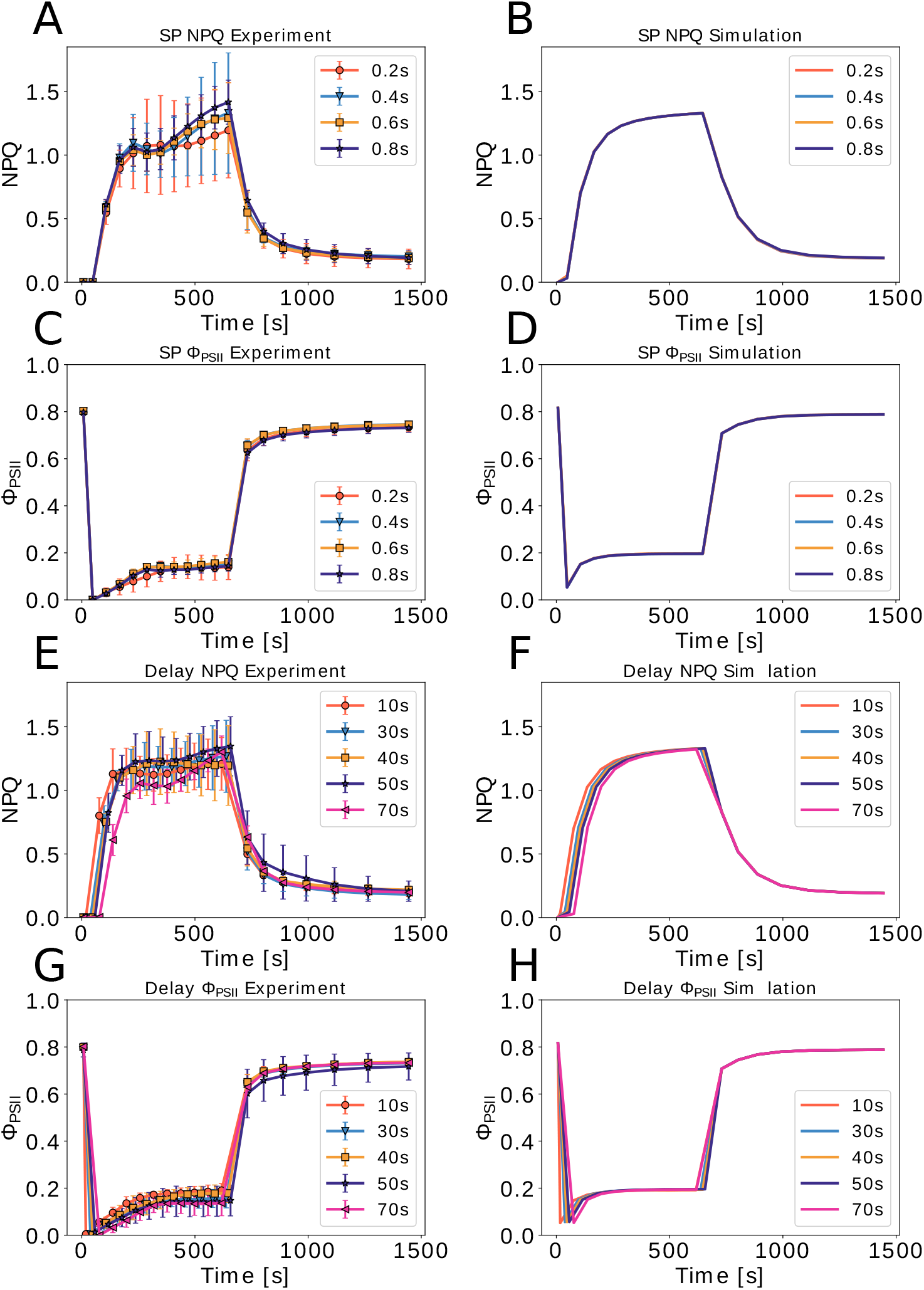
Comparison of the experimental and simulated NPQ and ΦPSII induction and relaxation. 8A to 8D: SP experiment with varying duration of the saturation pulse between 0.2 and 0.8 s. The delay was 40 s for all measurements. 8E to 8H: Delay experiment with varying delay to switch on the actinic light 10s, 30 s, 40 s, 50 s and 70 s after the F_m_ measurement. The duration of saturation pulse was 0.8 s for all measurements. Error bars indicate standard deviation (n=4).

**Fig. 9:**
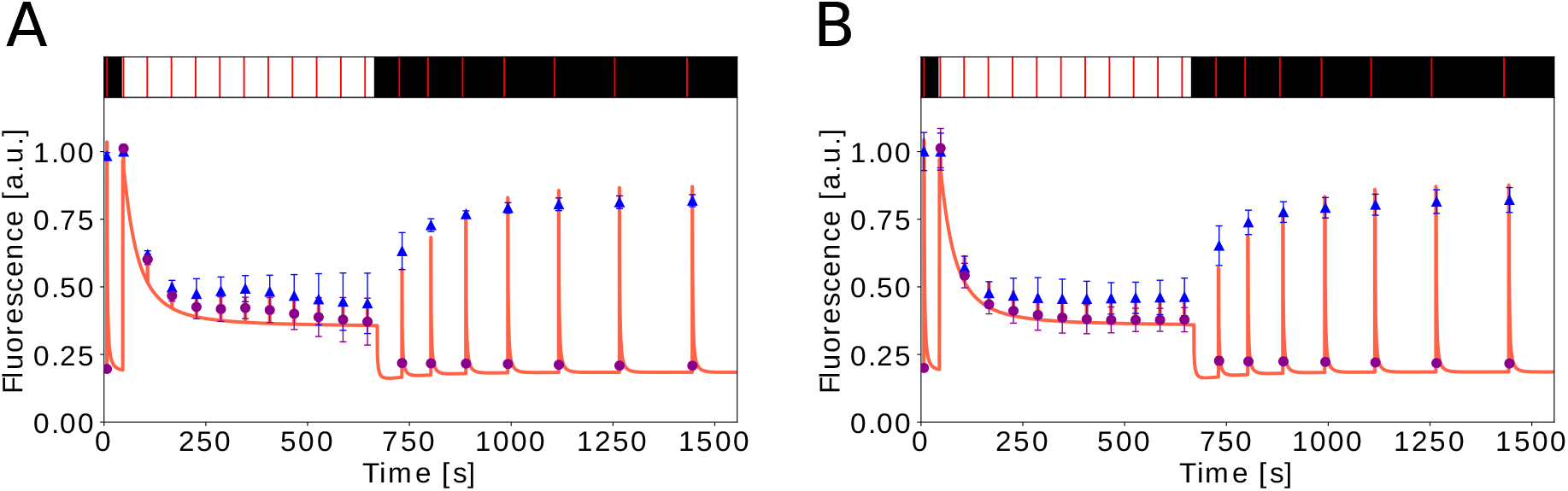
Comparison of the experimental data and simulated fluorescence traces. 9A: SP experiment with SP duration of 0.8 s and delay of 40 s. 9B: Delay experiment with a delay of 40 s and SP duration of 0.8 s. Blue symbols, mean values of F_m_ and F_m_’; purple symbols, mean values of F_o_ and F_s_; line, simulation. Error bars indicate standard deviation (n=4). Black and white bars above the fluorescence data show dark periods and actinic light illumination, respectively. Vertical red lines inside these bars indicate the time points of saturation pulse application.

In the SP experiment the model could reproduce some, yet not all features of the NPQ and Φ_PSII_ traces (Figs.8A to 8D). The duration of SP played only a minor role for both experimental determination and simulation of Φ_PSII_ (Figs. 8C and 8D). The model and experimental results are in good accordance for Φ_PSII_ (see section 3.4), with exception of somewhat faster kinetics of increase predicted by the model compared to the observation made in overnight dark-adapted plants. For NPQ the computational analysis deviated from the experiment during light induction whereas dark relaxation could be well reproduced (Figs. 8A and 8B). In addition, the experimental data, albeit showing large variations among the replicate plants, indicated that the NPQ value might be influenced by the duration of SP while the simulation did not show such influence (Fig. 8A)

These deviations may imply that the model parameters, despite the adjustments described in the Materials and Methods (2.1 Model used for simulations), are still suboptimal for the plants used for the experiments and/or that some mechanisms are not taken into account in the model. However, looking at the 0.8 s pulse data of the SP experiment and the 40 s delay data of the Delay experiment (Fig. 9), for which the same combination of SP duration (0.8 s) and delay (40 s) was used, it becomes evident that the plants in the latter experiment showed NPQ induction and fluorescence traces that were more similar to the simulation (Figs. 8 and 9). We do not know what factor(s) might have led to the different NPQ induction patterns in the two experiments. One possibility is that reactions in photosynthetic induction, which give rise to a highly variable PMST wave of the fluorescence induction curve (Papageorgiou and Govindjee, 1968a,b; Stirbet et al., 2014), contributed to the NPQ variations. It has been suggested that this slow phase of the fluorescence induction curve has multiple complex causes including the activation of the Calvin-Benson-Bassham cycle and state transition between the two photosystems induced by red and far-red light (Stirbet and Govindjee, 2016), both of which are not included in the simplified model used in this analysis. Since we used red AL in our experiments, the effect of blue light-induced chloroplast movement on NPQ induction (Cazzaniga et al., 2013) can be ruled out. Clearly, the quality (wavelength) of AL is another important information needed to interpret and simulate NPQ induction. Also the length of dark adaptation prior to the initial Fm measurement likely plays a role in the systematically different kinetics of Φ_PSII_ increase upon AL illumination found between the experiment and simulation (Fig. 8).

In the Delay experiment the expected changes in NPQ and Φ_PSII_ at the beginning and at the end of the AL phase were observed in the experiment as well as in the simulation (Figs.8E to 8H). These changes are only temporal shifts and do not indicate changes in NPQ activation or relaxation kinetics. Yet, as shown in the computational part, when SPs are not bound to the time of AL but fixed at given time points, the length of delay has an effect on NPQ (Figs. 3B and 4B). Hence, information about the time point of switching on the AL needs to be provided together with the information about the time point of the first SP. Overall, the comparisons between the experimental data and the model simulation shown in Figs. 8 and 9 support the main message of the simulation analysis presented in the sections 3.1 to 3.4, namely the importance to report all the details of the PAM experiment protocol for the sake of reproducibility and data interpretation as well as reusage of the experimental results. Furthermore, Figs. 8 and 9 show that the model (Matuszyńska et al., 2016) adequately reproduces the effects of small changes in technical PAM parameters and is able to give hints about which technical details are more important for the description of PAM protocols. When these details are known, deviations of simulations from experimental data can help us in the search for missing factors and mechanisms to improve the model.

## 4 Discussion

For decades now, PAM fluorescence measurements are widely used in plant research to assess e.g., plant health, genotypic variation, and effects of mutations. Due to their minimally invasive nature, fluorescence measurements provide a convenient method to assess photosynthetic dynamics *in vivo*. The power of this spectrometric technique lies in the connection between the yield of fluorescence and numerous intrinsic processes such as NPQ, making it a method of choice for many researchers to study oxygenic photosynthesis (Stirbet and Govindjee, 2011). Detailed understanding of the underlying mechanisms and regulating circuits of the photosynthetic machinery is essential for the improvement of agricultural and horticultural productivity and sustainability, such as better designing of greenhouses, and tailored plant breeding, as well as biotechnological exploitation. It is hence expected that the interest in applying fluorescence techniques will further increase in the future. In fact, more and more advanced technical devices and methods have been and are being invented that use fluorescence emission to obtain knowledge about the photosynthetic capacities of organisms, ranging from clip-on, leaf-level measurements, using for instance the MultispeQ device (Kuhlgert et al., 2016), to canopy-level measurements with the LIFT technique (Kolber et al. (2005) and proximal or remote sensing of solar-induced fluorescence (Aasen et al., 2019; Mohammed et al., 2019).

In parallel to the experimental efforts, numerous mathematical models simulating the dynamics of photosynthesis, often calibrated to PAM results, have been developed (for a review see Stirbet et al. (2014, 2019)). It was during one of such attempts to reproduce published results of PAM experiments *in silico*, where it came to our attention that a number of publications using the PAM technique do not provide all technical parameters of the experiments that are needed to simulate the results. Concerned about possible consequences of such omission, we carried out a systematic investigation of the effects of several key technical parameters on the output of PAM measurements. Using NPQ and ΦPSII, which are derived from the calculated fluorescence, we have quantified and visualised the differences between our computational experiments where such parameters have been varied. The exact time point of switching on and off the AL, combined with fixed time points of SPs, considerably affects the observed induction and relaxation kinetics of NPQ (Figs. 3 and 4). Because the kinetic information is often used to derive conclusions about photoacclimation and NPQ mechanisms, our simulations underline the importance of reporting the time points of AL and SPs applied in PAM experiments. We also examined the effects of the SP duration and the interval between those on fluorescence traces. While the former seems to have little direct influence on NPQ or PSII yield (Fig. 7), the latter had small but detectable effects on NPQ in low AL regimes (Fig. 6). Importantly, the combination of SP and AL intensities can influence the outcome of experiments substantially, as visualised by the steady-state NPQ landscape in Fig. 5. These mathematical simulations clearly demonstrate the importance of full disclosure of technical details. In fact, this conclusion holds true for many experimental studies, not only for PAM measurements. To exclude the possibility that it is the structure of the model that causes the observed differences, we repeated our simulations with another model of photosynthesis (Ebenhöh et al., 2014). Using this model, which has been developed with a focus on state transitions, another acclimation mechanism, still obtained similar results, suggesting that the simulated effects are not attributable to model structure (the analysis can be found in the same repository https://gitlab.com/qtb-hhu/fluopam). Moreover, we validated the *in silico* findings by conducting PAM experiments focusing on variations of the time point of switching on AL and the duration of the SP (Fig. 8). The predicted impact of these variations was largely in accordance with the experimental results. Although some level of reproducibility may be achieved between experiments by simply using the default settings of commercial instruments, this does not justify the omission of the details of experimental protocols. Especially, mathematical models and computational simulations have no default values, hence any missing information hinders further replication of studies. Without the necessary information, even a basic experiment cannot be properly simulated. The analysis presented above highlights the particular consequences of not providing detailed technical information about experimental protocols on the results of computational simulations of PAM Chl fluorescence traces of a photosynthetic organism. We hope that this article — our response to the ‘reproducibility crisis” — will reach a broad plant science community. We strongly encourage all readers, not only PAM users, to carefully and critically assess reporting practice to ensure independent replicability of experiments and to enable exploitation of results in the era of data science. As many have stated before us, the first step to increase reproducibility is to “increase the quality of protocol reporting” (Jessop-Fabre and Sonnenschein, 2019). We will not solve the problem of unreproducible and unreplicable research unless we provide and share all required information, even those seemingly minor and irrelevant ones. Finally, it is critical to reach an agreement on the minimal information that should be mandatorily given in the description of PAM experiments. This kind of agreements have already been made in the field of proteomics (MIAPE, Taylor et al. (2007)), next-generation sequencing (MINSEQE, http://fged.org/projects/minseqe/, accessed on: 16th April 2021) and microarray analysis (MIAME, Brazma et al. (2001)), in which databases are used extensively to analyse results of individual studies or selected sets of studies in form of Open Science. Also in the field of photosynthesis, PhotoSynQ is exploring a worldwide data sharing and analysis platform by creating low-cost devices and web-linked tools to collect and analyse data, exchange measurement protocols, or perform meta-analysis for registered users (https://www.photosynq.com/, accessed on: 6^th^ May 2021). A similar platform and database, where users can freely share experimental results and protocols, even including model algorithms for simulation, can be envisaged to promote the exploitation of PAM data and to accelerate knowledge exchange. For the realisation of such platforms, the data stored there must fulfill the minimal requirements to provide the information necessary for reproducing the experiments. Obviously, the information would comprise technical details of measurement protocols, as shown by the experiments and computational analyses of this article, as well as descriptions of plant materials, growth conditions, and treatments (Materials and Methods). Albeit outside the focus of this article, the importance of the latter information and the challenge to reproduce experimental results in plant research have been demonstrated by the joint experiments of 10 laboratories in the European AGRON-OMICS project (Massonnet et al., 2010). As was done in the aforementioned omics communities, a concrete and practical list of minimum information needs to be elaborated for PAM experiments by a consortium involving experimental and data scientists. By committing to such common standards and open databases, we will both contribute to and benefit from transparent, integrative, and interactive science in our research fields.

## Acknowledgements

This work was funded by the Deutsche Forschungsgemeinschaft (DFG, German Research Foundation) under Germany’s Excellence Strategy - EXC-2048/1 - project ID 390686111 (OE, AM, SM), Deutsche Forschungsgemeinschaft Research Grant MA 8103/1-1 (AM), Deutsche Forschungsgemeinschaft (DFG, project ID 391465903/GRK 2466) (TN), and Bundes Ministerium für Bildung und Forschung (BMBF, project ID 03SF0576A-B) (YN). We would like to thank all members of the Photosynthesis Task Force at the Institute of Quantitative and Theoretical Biology for the fortnightly meetings where the idea of this communication originates from. We also thank Roberto Bassi (University of Verona) for kindly providing the fluorescence emission spectrum of Chl *a*.

## Author contribution

Conceptualization: AM, OE; Data curation: TN; Formal analysis: TN, AM; Funding acquisition: SM, OE, AM; Investigation: TN, YN; Methodology: TN, YN, SM, OE, AM; Project administration: TN; Resources: SM, OE, AM; Software: TN, AM; Validation: TN, YN, SM, AM; Visualization: TN, AM; Writing - original draft: TN, AM; Writing - review & editing: TN, YN, OE, SM, AM.

## Data Availability Statement

The code to reproduce the computational analysis and the experimental data that support the findings are openly available on the GitLab repository https://gitlab.com/qtb-hhu/fluopam or can be requested from the authors.

